# Land-use change undermines the stability of avian functional diversity

**DOI:** 10.1101/2025.01.15.633202

**Authors:** Thomas L. Weeks, Patrick A. Walkden, David P. Edwards, Alexander C. Lees, Alexander L. Pigot, Andy Purvis, Joseph A. Tobias

**Affiliations:** Department of Life Sciences, Imperial College London, Silwood Park, Ascot, UK; Department of Life Sciences, Natural History Museum London, South Kensington, London UK; Department of Plant Sciences, University of Cambridge, Cambridge, UK; Department of Natural Sciences, Manchester Metropolitan University, Manchester, UK; Centre for Biodiversity and Environment Research, Department of Genetics, Evolution and Environment, University College London, London, UK

## Abstract

Land-use change causes widespread shifts in the composition and functional diversity (FD) of species assemblages, yet the impacts on ecosystem resilience remain uncertain. Stability of ecosystem functioning may increase after land-use change because the most sensitive species are removed, leaving more resilient survivors (Balmford, 1996; Clavel et al., 2010; McKinney & Lockwood, 1999). Alternatively, ecosystems may be destabilized if land-use change reduces functional redundancy, accentuating the ecological impacts of further species loss (Fonseca & Ganade, 2001; McCann, 2000). Current evidence is inconclusive, partly because trait data have not been available to quantify functional stability at sufficient scale. Here, we use morphological measurements of 3696 bird species to estimate shifts in functional redundancy following recent anthropogenic land-use change at 1281 sites worldwide, and then assess the sensitivity of these altered assemblages to future species loss. We find that the proportion of disturbance-tolerant species increases after land-use change, but that this does not increase stability. Instead, we show that functional redundancy is reduced and that further species loss will destabilize ecosystem function because relatively few additional extinctions lead to accelerated losses of FD, particularly in trophic groups delivering important ecological services such as seed dispersal and insect predation. Our analyses reveal that land-use change may have major undetected impacts on the stability of key ecological functions, hindering the capacity of natural ecosystems to absorb further declines in functionality caused by ongoing perturbations.

---

Anthropogenic land-use change is the primary driver of biodiversity decline and turnover (Jaureguiberry et al., 2023). It removes or changes local biodiversity through a variety of processes, including conversion of rich natural and semi-natural habitats to monocultures or urbanized settings with little vegetation (Foley et al., 2005; Grimm et al., 2008). These landscape transformations are a defining feature of the Anthropocene, leading to substantial shifts in the composition of local species assemblages (Allan et al., 2015; Etard et al., 2000; Flynn et al., 2009; Newbold et al., 2015). However, the impacts of these compositional changes on ecosystem function are difficult to measure or predict (Loreau et al., 2001; Thompson et al., 2018).

A standard approach to inferring changes to ecosystem function involves estimating the diversity of functional traits in species assemblages, based on strong evidence that species traits provide information about functional roles (Pigot et al., 2020; Cadotte et al., 2011; Petchey et al., 2004; Tilman et al., 1997). The correlation between traits and ecological processes has led to widespread use of functional diversity (FD) metrics to assess the impacts of land-use change on ecosystem function (e.g. Chapman et al., 2018; Flynn et al., 2009; La Sorte et al., 2018; Vandewalle et al., 2010; Villéger et al., 2008). Such analyses often conclude that land-use change has relatively minor effects on FD after accounting for species turnover (Edwards et al., 2013; Gorczynski & Beaudrot, 2021; Luck et al., 2013), and that high levels of functionality are therefore retained in human-modified landscapes (Magnago et al., 2015; De Coster et al., 2015). Nonetheless, the focus of most studies on overall trait diversity in assemblages has two main limitations. First, FD estimated for whole assemblages provides no information about the integrity of particular ecological functions, some of which are less resilient than others. Second, standard FD metrics reflect a snapshot in time and tell us nothing about the stability of ecosystem function in the face of further environmental perturbation (Fonseca & Ganade, 2001), suggesting that the long-term impacts of land-use change may be underestimated (Fig. 1a).

**Fig. 1.**
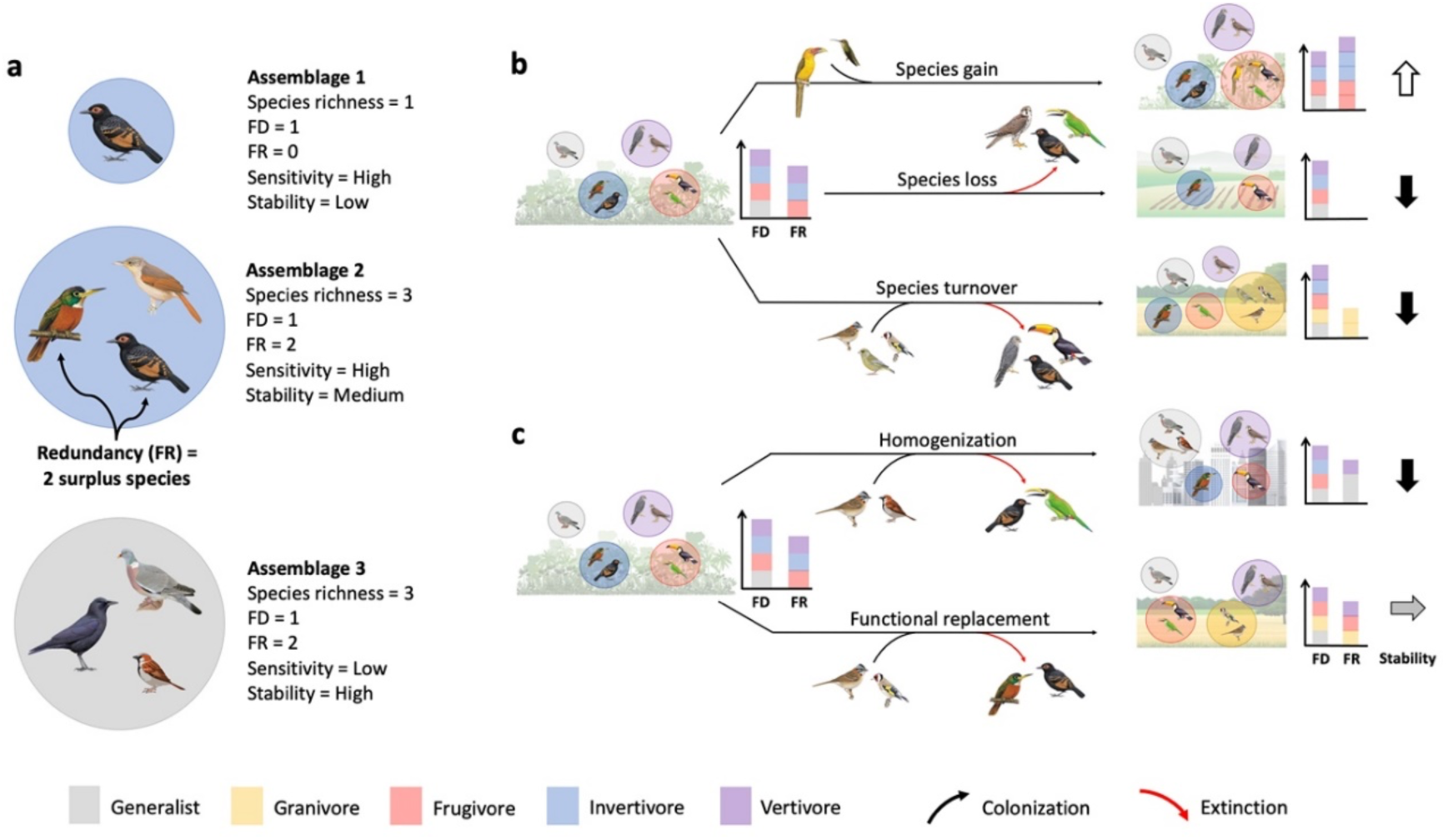
Impacts of land-use change on the structure, function and stability of species assemblages. (**a**) We define functional diversity (FD) in simplified form as functional group richness: the number of dietary guilds in the assemblage. Functional redundancy (FR) is the number of surplus species in each guild. Functional stability increases with the number of surplus species present. (**b,c**) Land-use transitions from primary vegetation drive changes in species composition of avian dietary guilds: generalists (grey), granivores (yellow), frugivores (red), invertivores (blue) & vertivores (purple). Different pathways of species extinction and colonization may have different impacts on FD, FR and stability, measured as Functional resistance (that is, the potential for assemblages to absorb further extinctions without declines in FD). Pathways in (**b**) show that FD and FR can fluctuate independently, and that stable or increasing FD can conceal decreases in both FR and Functional stability. Pathways in (**c**) show how FD and FR can fluctuate within functional groups without affecting assemblage-level metrics. In these cases, Functional stability can be impaired if species are disproportionately lost from more sensitive guilds.

Functional redundancy – and its flipside, functional uniqueness – are dimensions of FD that focus on the supply of species delivering each function. Redundancy metrics achieve this by estimating the number of co-occurring species with overlapping functionality (Biggs et al., 2020). If multiple species in an assemblage provide similar functions, surplus species are effectively functionally redundant (Lawton & Brown, 1994). In ecological terms, functional redundancy is a positive attribute (Eisenhauer et al., 2023) because surplus species increase resilience and stability, enabling continuity of ecological processes when conditions change (Fonseca & Ganade, 2001; McCann, 2000; Naeem, 1998; Naeem & Li, 1997). This ‘insurance effect’ means that species assemblages with high functional redundancy have greater functional resistance – that is, they are more stable because their overall functionality is maintained when species are lost from the assemblage (Oliver et al., 2015; Hillebrand et al., 2019).

Under random species loss, functional redundancy is equivalent to functional resistance. However, the effects of land-use change are non-random as some species with predictable characteristics are more extinction-prone and tend to be filtered from the new environment (Croci et al., 2008; Newbold et al., 2013, 2020). Moreover, these sensitive species are also distributed non-randomly, often clustering in distinct functional groups, which may undergo higher rates of local extinction, destabilizing the ecological processes they regulate (Burkle et al., 2013; Crooks & Soulé, 1999; Bregman et al., 2016). Indeed, if land-use change drives non-random species gains in some tolerant functional groups in parallel with species losses in more sensitive groups, the functional stability of an assemblage can be impaired despite no overall loss of functional redundancy (Fig. 1b). This may occur, for instance, when redundant ecological specialists are replaced by disturbance-tolerant or generalist species in anthropogenic habitats (Clavel et al., 2010; McKinney & Lockwood, 1999). We are left with a key conundrum: when humans alter the environment through land-use change, are the new species assemblages that form in anthropogenic landscapes more resistant to future shocks (because sensitive species are already lost and resilient species increase in abundance) or do they become more fragile and sensitive to further collapse?

To examine this question, we quantify the impacts of land-use change on functional trait diversity and redundancy of birds. Birds provide an opportunity to quantify the functional impacts of environmental change with unparalleled resolution because they have been surveyed intensively (Hughes et al., 2021), while comprehensive morphological trait data with well-established links to key ecological and trophic processes (Pigot et al., 2020) are now available for all bird species (Tobias et al., 2022). In total, we examined 3696 bird species across 1281 focal assemblages sampling land-use gradients from primary vegetation to urban habitats (Fig. 2a). We estimated the FD of each assemblage as the total volume (that is, functional richness) of the occupied trait space, quantified using probabilistic hypervolumes (Blonder et al., 2018; see Methods). When we compared 177 assemblages in primary vegetation – including 152 (86%) in forests and 25 (14%) in non-forest vegetation (mainly grasslands & shrublands) – with 1104 assemblages in nearby human-modified landscapes, we found that low-intensity human activity can drive slight but significant increases in FD, such as in disturbed primary vegetation (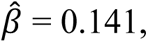 *P =* 0.043), or slight reductions in FD, as detected in mature secondary vegetation (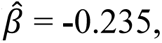 *P =* 0.046). However, substantial declines in FD were consistently observed across other, more heavily disturbed land-use types (Fig. 2b), most prominently in highly urbanized landscapes (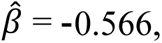 *P <* 0.001).

**Fig. 2.**
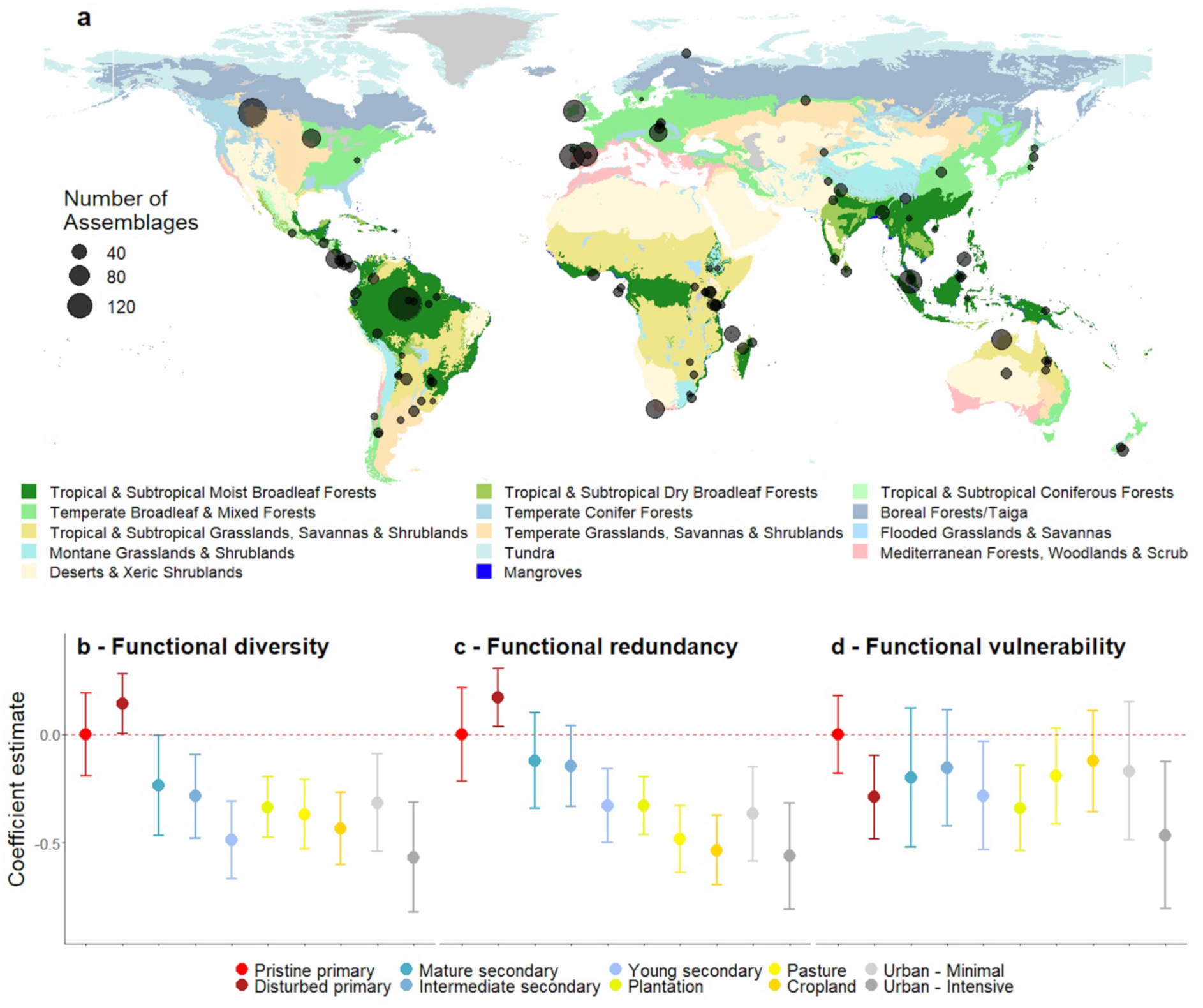
Sampling and impacts of land-use change on avian assemblages. (**a**) Circles show geographical location of 98 field surveys providing data for 1281 avian assemblages; circle size is proportional to the number of assemblages surveyed in each study. Colours indicate major biomes (Olson et al., 2001). Lower panel shows outputs from univariate mixed effects models assessing the impact of land-use change on three assemblage-level metrics: (**b**) functional diversity (FD; measured as functional richness), (**c**) functional redundancy (FR), and (**d**) functional vulnerability (FV). All metrics are compared to a pristine primary vegation baseline (including forests, grasslands, shrublands and wetlands; dashed red line). FV is calculated using the Pearson’s correlation coefficient between species-level redundancy and trait-based sensitivity scores (see Methods). Results shown are coefficient estimates and 95% confidence intervals. Response variables were squareroot transformed prior to analysis and then scaled by their standard deviation to aid comparison (the transformation step was not possible for FV). Note that -ve FV implies greater resistance to further loss of FD.

## Functional redundancy

To evaluate these patterns in FD from the perspective of ecosystem stability, we calculated functional redundancy for each assemblage as the amount of shared niche overlap between co-occurring species. To do this, we constructed a hypervolume using comprehensive trait measurements of all species in the assemblage (Tobias et al., 2022), then estimated the position and extent of niche-space for each species based on intraspecific variation within the hypervolume. This approach revealed that, in addition to its effects on FD, land-use change also alters assemblage-level functional redundancy (Fig. 2c), roughly matching patterns reported from plant communities (Laliberté et al., 2010). Specifically, we show that trait redundancy initially increases after the switch from pristine to disturbed primary vegetation (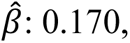 *P =* 0.012), with minor, non-significant declines in both mature (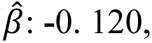 *P =* 0.288) and intermediate-age (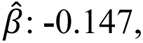 *P =* 0.122) secondary vegetation. However, more intensive land-use types showed significant decreases in redundancy, with particularly sharp declines in cropland (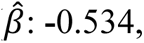 *P* < 0.001) and intensively urbanized landscapes (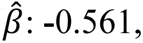 *P* < 0.001). Our findings were similar whether we calculated intraspecific variation using a standard kernel density estimator (Carmona et al., 2016; see Methods) or based on direct measurements of multiple individuals per species (Supplementary Fig. 1).

When we modelled change in FD and functional redundancy within four different functional guilds, responses to land-use change were highly divergent. In dietary generalists and granivores, FD and redundancy remained constant or increased in agricultural and urban landscapes, reflecting an influx of open-country and urban-tolerant species, some with distinctive traits (Extended Data Fig. 1). In contrast, FD and redundancy of frugivores (involved in seed dispersal) and invertivores (with roles in controlling insect populations) declined sharply (Extended Data Fig. 1). These findings indicate that whole-assemblage FD should be treated with caution because it averages across multiple ecological processes with widely diverging sensitivity to land-use change (see Supplementary Information). Increased diversity in disturbance-tolerant guilds can obscure substantial declines in disturbance-sensitive guilds, masking a diminished capacity of assemblages to maintain important ecosystem functions.

## Functional vulnerability

Although these functional redundancy patterns imply that land-use change can limit the capacity of anthropogenic ecosystems to withstand further species losses, the link between functional redundancy and stability is not clear-cut. Assemblages with low redundancy can be stable if the remaining species are well-adapted to human-modified landscapes. Moreover, ecological functions can be unstable even in highly redundant assemblages if many species are densely packed into only a few functional groups leaving other areas of trait-space under-represented (Mouillot et al., 2014). In such a scenario, the delivery of rarer functions can be unstable if the species responsible are disproportionately sensitive to land-use change or persist at small population sizes (Curtis et al., 2021; see Methods).

To examine the question of stability more closely, we devised two metrics of functional vulnerability (FV) which incorporated the amount of unique function provided by species, along with their likely sensitivity to anthropogenic pressures. Specifically, we calculated a species-level redundancy value based on the relative contribution of each species to total assemblage functional redundancy and a sensitivity score for each species based on their sensitivity to disturbance. We estimated sensitivity based on species-specific response traits or population size (rarity), representing the likelihood that each species would undergo local extinction in response to further environmental change (see Methods). To generate FV values, we calculated the covariance between the sensitivity scores for all species occurring in the assemblage and their functional redundancy. High FV values indicate a negative covariance between sensitivity and redundancy implying that species with heightened extinction risk also provide a large proportion of unique function.

We found that land-use change drives consistent declines in FV: all anthropogenic land-use types other than secondary vegetation underwent significant decreases in at least one vulnerability metric. The declines were similar whether species’ sensitivity to disturbance was estimated as a function of response traits (trait-based FV; Fig. 2d) or abundance (rarity-based FV; Extended Data Fig. 2). These findings support the hypothesis that land-use change removes the most sensitive species, resulting in lower assemblage vulnerability because most of the species surviving in and colonizing anthropogenic landscapes tend to be less prone to extinction (Betts et al., 2019; McKinney & Lockwood, 1999). However, FV metrics may over-estimate the future stability of ecosystems if they do not consider how redundancy is distributed across the species assemblage as a whole.

## Future extinctions and the stability of ecosystem function

Our finding that both functional redundancy and FV decline after land-use change suggests that human impacts may have opposing effects on ecosystem stability. Each ecological function may be delivered by a reduced set of species, yet these species have lower extinction risk because they are more tolerant of further anthropogenic pressures. To disentangle these effects, we first estimated future ecological stability in the form of functional resistance to simulated extinctions. We calculated two functional resistance values for each assemblage by removing species in order of sensitivity (high to low) using the same trait-based and rarity-based sensitivity scores used to calculate FV values (Fig. 3a; see Methods). This allowed us to track the rate at which FD declined when species were sequentially removed from the assemblage (Fig 3b; Extended Data Fig. 3; Oliver et al., 2015; Hillebrand et al., 2019) and then to test whether these estimates of functional resistance are best explained by redundancy or FV.

**Fig. 3.**
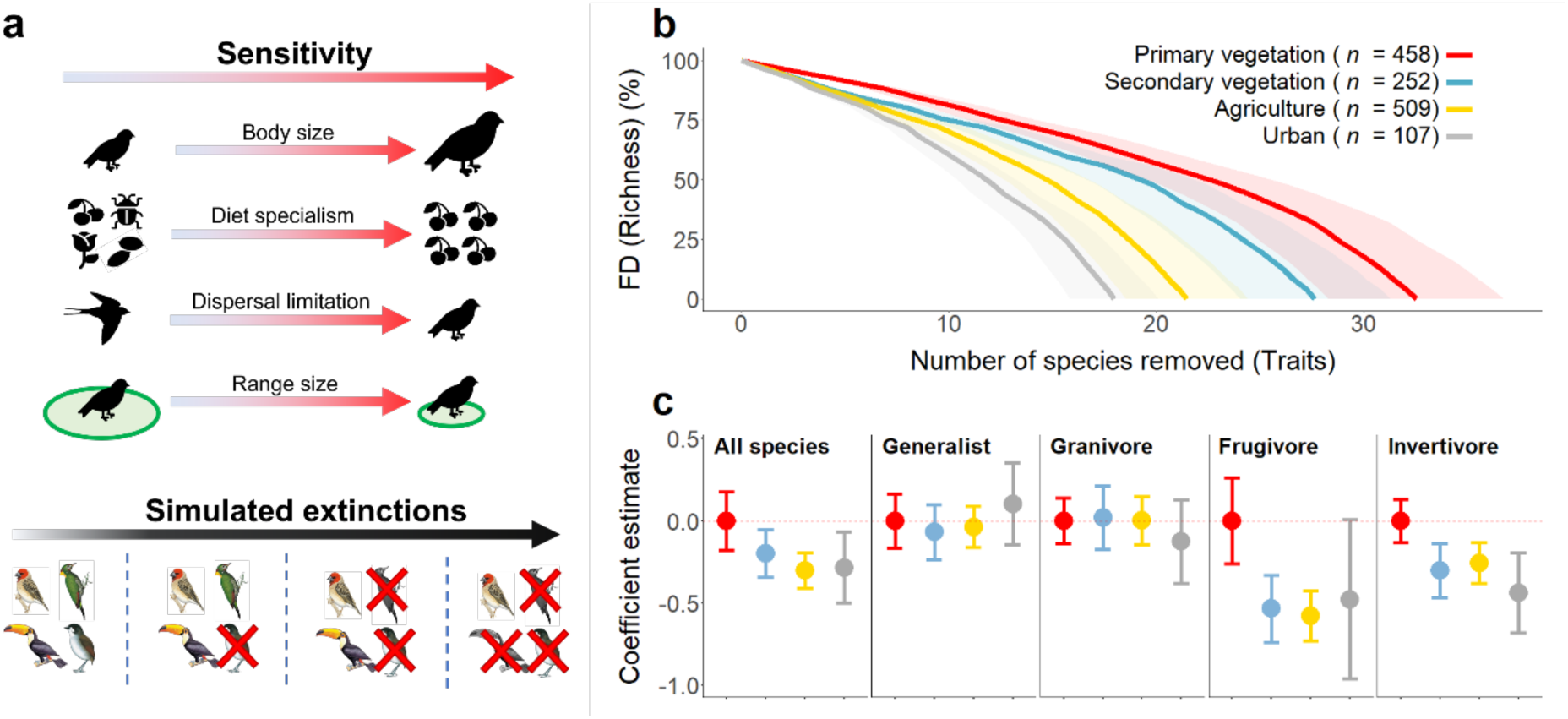
Land-use change reduces functional stability. **a**, Procedure used to quantify functional stability of assemblages (*n* = 1281). As a first step, all species in an assemblage are ranked by extinction risk based on four response traits (see Methods), then simulated extinction curves are generated by removing species sequentially in order of their sensitivity score (high-to-low). The impact of species loss can then be quantified by calculating functional trait diversity (FD) as a proportion of the starting FD before any species were removed, providing an index of stability (functional resistance). Using this approach, we plot average extinction curves (**b**) for each land-use type predicted using a cubic smooth spline algorithm. To aid visualization, we use the total number of species removed rather than proportional data. The impacts of land-use change on functional resistance in different trophic groups (**c**) can be visualised by calculating predicted change in functional resistance as the area under the extinction curve (AUC; see Extended Data Fig. 3); dashed red line indicates the predicted functional resistance for all primary vegetation types (forests, shrublands and grasslands). Metrics are calculated and compared across five subsets: all species, trophic generalists, and three key functional guilds. Results shown are coefficient estimates from five separate linear mixed-effects models with 95% confidence intervals.

Using bivariate models, we found that assemblage-level redundancy is positively correlated with functional resistance (trait-based 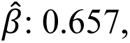 *P* < 0.001; rarity-based 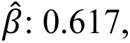 *P* < 0.001; Extended Data Fig. 4). Conversely, the effect of FV on resistance is significantly negative (trait-based 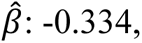 *P* < 0.001; rarity-based 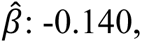 *P* < 0.001), indicating that functional resistance is undermined when extinction-prone species provide unique functions that cannot be supplied by other species in an assemblage. This finding suggests that realistic extinction processes – which are likely to target the most extinction-prone or rarest species – may initially cause FD to decline in assemblages with high FV. This may occur, for example, in habitats supporting bird assemblages with a higher proportion of rare and morphologically unique species, such as primary vegetation types (Fig. 2d; Extended Data Fig. 2). Nonetheless, the relatively weak effect of FV suggests that any declines in assemblage-level vulnerability may be counteracted by the stronger effect of declining redundancy across the entire assemblage.

To examine this hypothesis further, we assessed how land-use change influences the functional stability of assemblages undergoing future anthropogenic extinctions. We found that functional resistance for whole assemblages follows a similar pattern to functional redundancy, with substantial declines occurring in human-modified landscapes, particularly in agricultural and urban settings (Fig. 3c). Finally, we ran sensitivity analyses with an alternative functional resistance metric: the half-life (t_1/2_) of each extinction curve, defined as the proportion of species that need to be removed for FD to decline by 50% (Fonseca & Ganade 2001; Extended Data Fig. 3). The results were similar, suggesting that functional stability consistently declines with increasing land-use intensity (Supplementary Fig. 2).

## Ecological and geographical factors regulating functional stability

Functional stability calculated at the assemblage level reflects an average across different trophic groups with varying responses to land-use change (Newbold et al., 2013; 2020). Accordingly, we find that functional resistance in assemblages of dietary generalists and granivores tends to remain stable or even to increase across human-modified landscapes, whereas the ability to maintain a diversity of functions becomes highly unstable in more sensitive guilds, including frugivores and invertivores (Fig. 3c; Extended Data Fig. 1). Declines in ecosystem stability after land-use change are therefore unevenly distributed, with the largest losses concentrated in key trophic guilds mediating ecological services, such as seed dispersal and pest control, which are vulnerable to future collapse in human-modified landscapes. The sensitivity of these guilds to land-use change is consistent with previous studies showing that frugivorous and insectivorous birds are particularly susceptible to local extirpation in disturbed tropical forests (Şekercioğlu et al., 2002; Bregman et al., 2016).

Indeed, the response of tropical seed dispersers and insect predators to land-use change drives a more general pattern of accentuated declines in FD and redundancy at lower latitudes, where many bird species have high sensitivity to habitat loss and fragmentation (Bregman et al., 2014; Weeks et al., 2023; Extended Data Fig. 5). In such cases, the relatively higher redundancy associated with intact tropical assemblages does not always result in higher functional resistance (Supplementary Fig. 6).

## Implications for ecosystem stability and land-use management

The widespread decline we show in avian FD is consistent with numerous studies reporting reduced FD in response to urbanization, agricultural expansion, and land-use intensification (Allan et al., 2015; Etard et al., 2022; Fischer et al., 2007; Flynn et al., 2009; Sol et al., 2020). This outcome reflects the loss of species maladapted to highly modified environments, including ecological specialists occupying unique regions of functional trait space (La Sorte et al., 2018; Oliveira Hagen et al., 2017). We also detect substantial reductions in trait redundancy and assemblage-level functional vulnerability, suggesting that human-modified assemblages are dominated by fewer, typically generalist species as landscapes become more intensively transformed (Clavel et al., 2010). The lower functional vulnerability of post-disturbance assemblages implies that they are more resilient, perhaps because extinction filters have removed the most sensitive species (Balmford, 1996; Betts et al., 2019). However, the results of simulated extinctions suggest the opposite (Fig. 3). Instead, as redundancy declines, the insurance effect provided by the rich diversity of undisturbed bird assemblages is eroded, accentuating the adverse impacts of further species losses (Fonseca & Ganade, 2001; Oliver et al., 2015). In other words, the minor positive effects of land-use change on FV are outweighed by reductions in trait redundancy, undermining functional stability by leaving ecosystems susceptible to much larger declines in functionality if further species are lost.

Our results indicate that disturbance-sensitive species in pristine habitats are typically functionally unique, highlighting the role of intact ecosystems as safe harbours for rare, functionally distinct, and extinction-prone species (Barlow et al., 2007; Mouillot et al., 2013; Weeks et al., 2022a). It may seem logical to conclude that ecosystem functionality is least stable in natural primary vegetation where so many disturbance-sensitive species are important to ecological function (Díaz et al., 2013). However, we show that greater instability arises from widespread reductions in trait redundancy occurring throughout the entire assemblage in moderately to heavily disturbed environments (Extended Data Fig. 4), consistent with theoretical predictions and experimental evidence (McLean et al., 2019; Nunes et al., 2021; Tilman et al., 2006). Undisturbed and moderately disturbed habitats support much higher levels of redundancy throughout the entire assemblage, thereby promoting functional stability. Importantly, well-developed secondary vegetation and lightly disturbed habitats have similar levels of redundancy to those found in pristine primary vegetation, highlighting the importance of retaining and restoring semi-natural and disturbed vegetation to boost the resilience of ecosystems functions (Edwards et al., 2014a).

## Conclusions

Our analyses reveal that land-use change drives pervasive declines in functional resistance and stability in bird assemblages. The impacts are most severe in heavily modified environments and concentrated in key ecological groups with prominent roles in seed dispersal (frugivores) and pest control (invertivores). The consistent global pattern detected in birds confirms and extends the findings of local-scale studies showing reduced functional resistance in invertebrate assemblages (Clapcott et al., 2012; De Vries et al., 2021). An important implication of these findings is that standard approaches to estimating the effects of land-use change on ecosystem function may underestimate longer-term impacts. Specifically, they may suggest that species assemblages in human-modified landscapes are more resilient, whereas a more detailed appraisal using trait hypervolumes reveals they are actually more fragile and primed for further declines in functionality if biodiversity losses continue unchecked (Leclère et al., 2020). Conservation efforts should therefore focus on boosting functional redundancy of species assemblages through habitat management to reduce the risk of future ecological collapse.

## Methods

### Survey data

To collate bird survey data, we started from a baseline of 145 studies comparing avian diversity at different stages of land-use change in the PREDICTS database (Hudson et al., 2014). We removed 39 surveys because they duplicated a single site (*n* = 1), lacked abundance data (*n* = 19), or were otherwise incomplete, usually because the sampling was limited to particular guilds or methods, such as camera traps (see Supplementary Information). To improve sampling in species-rich regions, we integrated data from intensive surveys in the Amazon (Nunes et al., 2022) and Bornean rainforests (Edwards et al., 2014b). Data were adjusted to align with the same nested sampling design of PREDICTS (see Supplementary Information).

To classify land-use types at each study site, we used PREDICTS data to estimate the predominant type and stage of vegetation, and the intensity of human use. First, we assigned sites to one of six vegetation classes (primary vegetation, secondary vegetation, plantation forests, pasture, cropland, urban). We then classified lightly or intensively used primary vegetation sites as disturbed primary vegetation, and minimal-use primary vegetation sites as pristine-primary vegetation (a proxy for undisturbed natural vegetation). Based on previous analyses showing reduced avian FD in intensely urbanized areas (Sol et al., 2020), we also split minimal-use urban sites from sites with more intensive urbanization. Finally, to account for the effects of vegetation structure at different successional stages (Drapeau et al., 2000), we partitioned secondary vegetation according to age class (Mature, Intermediate, Young, Indeterminate; see Supplementary Information). Indeterminate age secondary vegetation was removed from our dataset.

Most survey data in the PREDICTS database is organised as a hierarchy, with survey sites (assemblages) nested within temporal or geographical blocks, and blocks nested within overall study sites. To fit this format, we restructured additional data by condensing 17 independent studies into seven studies with 2–5 blocks each (see Supplementary Information). Survey techniques and sampling effort vary across studies, which there differ in the extent of area sampled at each site. In some studies, each point-count is considered a single site which can cause problems for our analyses because the full species assemblage is poorly represented and neighbouring sites may be very close together. To improve sampling depth and to minimize the risk of resampling (pseudo-replication), we aggregated different sampling points within the same study-block into single sites (assemblages) when the points were close together, as well as in similar land-use types with similar use intensity.

Our final dataset consisted of 98 studies across six continents (Fig. 2a; Supplementary Table 3) and a total of 1281 avian assemblages in 10 distinct land-use types: pristine primary vegetation (*n* = 177), disturbed primary vegetation (*n* = 281), mature secondary vegetation (*n* = 44), intermediate age secondary vegetation (*n* = 77), young secondary vegetation (*n* = 86), plantation forest (*n* = 218), pasture (*n* = 184), cropland (*n* = 107), and urban, including both minimal-use (*n* = 46) and intense-use (*n* = 61) urban landscapes.

### Functional trait data

Morphological functional traits were obtained for all 3696 species recorded in our study assemblages. Species-mean morphological trait values were compiled from the AVONET database (Tobias et al., 2022) for seven biometric traits: beak length (culmen), beak length (tip-to-nares distance), beak depth, beak width, tail length, tarsus length and wing length. These traits have been shown to predict a range of key ecological niche axes, including diet and foraging strategy (Pigot et al., 2020). We also included data on Hand-wing Index (HWI), a metric of wing elongation that predicts dispersal distance in birds (Weeks et al., 2022b), and thus widely used as a proxy for dispersal ability (Sheard et al., 2020; White, 2016). Mean values for all morphometric traits were calculated from an average of 11 individuals per species (41515 individual birds measured in total).

Some taxa reported in survey data were impossible to assign directly to species because they were only identified to genus level (see Supplementary Information). Deleting these taxa would result in missing data, which can reduce the accuracy of FD estimates (Pakeman, 2014). Instead, we created pseudo-species representative of the genus. Given that avian life history and morphological traits tend to be highly conserved within genera (Tobias et al., 2022), we assigned trait values to pseudo-species by averaging the trait values of all congeners potentially occurring at the locality. To generate trait data for averaging, we used geographical range polygons from BirdLife International (2021) to provide a list of all members of the focal genus with geographical distributions overlapping the site location. We synthesized data for 73 pseudo-species in 133 of our 1281 study assemblages.

Avian morphological traits are often intercorrelated because of an underlying association with body size (Pigot et al., 2020; Extended Data Fig. 6). Accordingly, all traits in our dataset were strongly correlated with the body size axis apart from HWI (*R* = 0.22). Following previous studies (Bregman et al., 2016; Trisos et al., 2014), we removed the association with body size through a two-step principal component analysis (PCA) which reduced our seven linear morphometric traits into three niche axes related to ecological functions (Extended Data Fig. 7). We performed two separate PCAs on trophic traits (related to beak morphology) and locomotory traits using all species in our dataset. In both cases, the first principal component (PC) is strongly correlated with body size, so we used the second PC to represent the dominant axis of variation (Extended Data Fig. 8). We then performed a separate PCA on the first PC scores from both the trophic and locomotory PCAs. The first PC was taken to represent the body size axis. We supplement these three derived trait axes with a fourth morphological trait axis consisting of the log-transformed HWI (related to dispersal ability).

### Dietary data

Birds mediate a wide range of ecological processes and services depending on their trophic interactions, including seed dispersal by frugivores and pest control by invertivores (Şekercioğlu, 2006; Whelan et al., 2015). The morphological trait dataset we use in this study has previously been shown to strongly predict avian diets and behavioural foraging strategies (Pigot et al., 2020). However, the connection between morphology and diet is noisy and weaker in some taxonomic groups (Bright et al., 2016; Felice et al., 2019). Therefore, we included standard diet classifications in functional metric calculations. We used published estimates of the proportion of species diets across nine major resource types: herbivore (aquatic), herbivore (terrestrial), nectarivore, granivore, frugivore, invertivore, vertivore (aquatic), vertivore (terrestrial), scavenger. This data was extracted from Pigot et al. (2020) and is primarily based on the EltonTraits dataset (Wilman et al., 2014) with extensive corrections, updates and reorganization based on recent literature. We also extracted trophic niche data from AVONET (Tobias et al., 2022) which identifies the primary food type exploited by each species obtaining at least 60% of its diet from a single food type. Species obtaining resources more equally across different food types were classed as Omnivores (Burin et al., 2016; Pigot et al., 2020).

### Calculation of functional diversity and redundancy

To calculate functional metrics, we used species’ diet proportion data across nine major resource types and their position on our four derived morphological trait-axes (Locomotory, Trophic, Dispersal and Size) to create Trait Probability Densities (TPDs) using the *TPD* package in R (Carmona et al., 2019). This approach uses species mean trait-values and intraspecific variation to calculate probabilistic hypervolumes that predict the potential position of species in functional trait space (representing a Hutchinsonian niche; Blonder et al., 2018). We then estimated FD as the total volume occupied in trait space (that is, functional richness) and calculated functional redundancy as the proportion of this volume that is shared by multiple species (see Supplementary Information; Carmona et al., 2019).

To create TPDs, we first calculated distance matrices using the package *gawdis* in R which is designed to handle grouped and proportional data (de Bello et al., 2021). We calculated a distance matrix for each of our 98 studies using diet and morphological data for all species present in each study. Following methods described by Legendre & Legendre (1998) we back-transformed our distance matrices into three-dimensional coordinates representing the relative position of each species in functional trait space (see Supplementary Information).

In addition to the species-mean position in a three-dimensional trait space, calculation of TPDs requires an estimate of intraspecific variation along each dimension. This estimate is used to calculate a kernel probability density around each species coordinate. Following previous studies, we estimated intraspecific variation for all species using a plug-in kernel density bandwidth estimator, which implements the Hpi.diag function from the *ks* package (Carmona et al., 2016, 2021; Duong, 2007). This method estimates a separate bandwidth for each dimension of trait space which represents intraspecific variation within each niche axis. We supplied the three-dimensional coordinates, derived from our distance matrix, and the square root of intraspecific variation estimates to the *TPDMeans* function to our study-level TPDs, which we then reduced into smaller site-level TPD subsets that reflect the position of all species present in each individual site. To assess the sensitivity of our results to kernel density methods (Carmona et al., 2016), we re-ran the analyses after estimating intraspecific variation from direct measurements of 18183 individual birds: a mean sample size of 5 individuals per species (Tobias et al., 2022).

To analyse the effect of land-use change on specific ecosystem functions, we recreated our TPDs for four species subsets to assess how functional trait structure changed within different functional guilds. Calculation of TPDs requires a minimum of four species to be present in any particular assemblage so sample sizes vary: all species (*n* = 1281), generalists (*n* = 1256), granivores (*n* = 1051), frugivores (*n* = 617) & invertivores (*n* = 1264). Finally, we calculated the FD and functional redundancy of each TPD using inbuilt functions in the TPD package (Carmona et al., 2019).

To address whether our estimation of intraspecific variation affected our conclusions, we recreated our TPDs using intraspecific variation calculated from multiple measurements of conspecific individuals extracted from AVONET (Tobias et al., 2022; see Supplementary Information). The results of sensitivity analyses based on these revised TPDs were similar, indicating the same general patterns of decline in FD or redundancy with land-use change (Supplementary Fig. 1).

### Calculation of species-sensitivity

Within each assemblage we calculated sensitivity scores for each species which are later used in our calculation of functional vulnerability (FV) and functional resistance metrics. We calculated two separate sensitivity scores which represent the likelihood that the species would be removed from the assemblage under distinct scenarios i) trait-based (species sensitivity driven by response traits), and ii) rarity-based (species sensitivity driven by abundance: low to high). Our trait-based sensitivity score was based on four key response traits associated with extinction risk: geographical range size, body size, diet specialism and dispersal limitation (Gaston & Blackburn, 1995; Owens & Bennett, 2000; see Supplementary Information). To calculate our rarity-based sensitivity scores, we extracted the inverse (that is, negative) abundance of each species within the assemblage based on the assumption that rarer species are more likely to be removed from an environment by population fluctuations (Lande 1993; Curtis et al., 2021). All sensitivity scores were scaled by their standard deviation and centred to have a mean of 0.

### Functional vulnerability

Previous studies have associated functional stability with the distribution of unique functional traits (Zhang & Zang, 2021), and linked assemblage vulnerability to the variation in sensitivity of species traits (Weeks et al., 2022a). We devised a new metric of functional vulnerability (FV) to encapsulate both these concepts by quantifying the relationship between the redundancy and sensitivity of the species in the assemblage.

To assess the distribution of sensitivity in relation to redundancy in species assemblages, we quantified the amount of redundancy provided by each species. Species-redundancy was estimated as the change in assemblage-redundancy after removal of the focal species. To calculate this value, we separately removed each species from the full assemblage and recalculated the assemblage-redundancy. We then calculated two functional vulnerability (FV) scores as the covariance between species-redundancy and either trait-based or rarity-based sensitivity scores using Pearson’s correlation coefficient. We reverse the direction of the covariance so that a high positive correlation indicates that the most sensitive species in the assemblage tend to be the least redundant (that is, most unique).

### Calculating functional resistance

We ran species extinction simulations and quantified the pace of FD declines with sequential species removals. A species’ sensitivity to environmental change is strongly influenced by certain functional response traits (Suding et al., 2008; Newbold et al., 2013, 2014). By extension, the likelihood of a species being removed from an assemblage is non-uniform and is instead likely a function of certain trait combinations. In addition, species abundances are often a strong indicator of local extinction risk as rarer species are typically more sensitive to environmental perturbations (Davies et al., 2004; Şekercioğlu et al., 2004). Therefore, for each assemblage we ran two extinction simulation scenarios by sequentially removing each species according to both sensitivity scores (trait-based and rarity-based).

Following previous methods, we plotted extinction curves to quantify how FD declines as the proportion of species occurring in the original assemblage is reduced to 0 (Carmona et al., 2016; Galland et al., 2020; Leitão et al., 2016; Sasaki et al., 2017). Using a standard approach, we then measured the area under the extinction curve (AUC) as an estimate of functional resistance (Leitão et al., 2016; Sasaki et al., 2017). If an assemblage maintains constant FD when species are removed, the AUC remains large, indicating high levels of functional resistance. Conversely, when extinctions drive declines in FD, the AUC decreases (Extended Data Fig. 3). As we are specifically interested in quantifying the pace of FD decline, we standardized the extinction curves by scaling the FD values between 1 and 0, with values close to 1 reflecting FD scores close to the starting value. This standardization prevents the magnitude of the FD value from affecting the size of the AUC values (Galland et al., 2020). This, variation in AUC values reflects variation in the shape of the extinction curve only.

Similar to our FD and redundancy analyses we calculated our functional resistance metrics across five different species subsets. We focused on the entire assemblage in addition to four key dietary-guilds which had been sufficiently sampled across all land-use types: all species (*n* = 1281), generalists (*n* = 1256), granivores (*n* = 1051), frugivores (*n* = 617) & invertivores (*n* = 1264). Sample sizes vary between functional guilds as TPDs cannot be created when three or fewer members of the guild are recorded in a given study. Therefore, in many studies we are unable to calculate initial FD estimates, meaning we cannot calculate resistance metrics for every guild across all study-sites.

Species in the same assemblage can have the same trait- and rarity-based sensitivity scores if they share a similar combination of response traits or are equally abundant. Therefore, we repeated our analyses 100 times and randomized the order which we removed species with similar sensitivity scores with each iteration. For each iteration we calculated an AUC value and took the mean across all values as our final functional resistance estimate.

One drawback of the AUC approach is its sensitivity to the order in which species are lost from an assemblage. For example, when a single morphologically unique species is lost before other more redundant species, this causes a steep initial decline in FD. Therefore, we also calculated functional resistance as the half-life (t_1/2_) of each extinction curve, defined as the proportion of species that need to be removed for FD to decline by 50% (Fonseca & Ganade, 2001; Extended Data Fig. 3). When we ran sensitivity analyses using this metric, we found that results were similar, supporting our conclusions (see Supplementary Information).

### Statistical analyses

To assess the impact of land-use on FD and functional redundancy, we first performed a set of fourteen univariate linear mixed effects models with land-use as the sole predictor variable. This set of analyses consisted of one model assessing the effects of land-use on FD and another model assessing effects on redundancy. These models were conducted across all species and separately across four different data subsetsrelated to dietary guild (generalists, granivores, frugivores and invertivores), and two subsets related to climatic region (tropics and non-tropical). We also ran two additional sets of three univariate models, which assessed whether trait-based or rarity-based FV were affected by land-use change. In both cases these two models were run across three data subsets: i) all studies, ii) tropical studies, iii) non-tropical studies. Rarity-based FV was unusually high for three sites as they contained extremely high variation in abundance within the site (that is, some species had very low abundance while others had extremely high abundance). Therefore, we removed these sites from rarity-based FV analyses.

In all models, we added study as a random effect to account for among-study differences in sampling methods. We also included study-block as a random-effect to account for within-study differences in the spatial and temporal structure of the sites. Due to the hierarchical structure of our data, it is difficult to incorporate covariance structures that account for spatial autocorrelation between local sites in the same study into our global models. However, following Newbold et al. (2015), we assessed spatial autocorrelation within studies and study-blocks separately using Moran’s I and found that it did not affect our results (see Supplementary Figs 3-5). Land-use was split into 10 distinct categories: pristine-primary vegetation, disturbed primary vegetation, mature secondary vegetation, intermediate age secondary vegetation, young secondary vegetation, plantation forests, pasture, cropland, minimal urban, and intense urban. Models were interpreted by comparing the change in response variable compared to Pristine primary vegetation.

We ran a third set of bivariate linear models to assess the proximal drivers of functional resistance calculated under both of our species-loss scenarios (trait-based and rarity-based). For each scenario, we ran a linear model with functional redundancy and FV as predictors of functional resistance with no associated random effects. For our trait-based and rarity-based scenarios, we used trait-based and rarity-based FV scores, respectively.

Finally, to address how land-use change ultimately impacts the functional resistance of bird assemblages, we ran two additional univariate mixed effects models. These analyses modelled the effects of land-use change on functional resistance of each assemblage under both species-loss scenarios (trait-based and rarity-based). We included study and study-block as random effects, in line with our first set of univariate mixed-effects models. We also split land-use into four categories: i) primary vegetation, ii) secondary vegetation, iii) agriculture (including plantation forests), and iv) urban. Models were interpreted by comparing the change in estimated effect size for each land-use type using primary vegetation as our reference category.

## Acknowledgements

We thank Sara Contu for assistance with methods and accessing survey data. TLW was funded by the Natural Environment Research Council studentship through the Science and Solutions for a Changing Planet Doctoral Training Programme.

## Author contributions

TLW and JAT conceived and developed the study, with input from AP and ALP. TLW integrated datasets and ran all analyses with support from PAW and AP. DPE and ACL provided additional datasets. TLW wrote the first version of the manuscript and designed all figures with input from JAT. All authors contributed to subsequent drafts and gave final permission for publication.

## Data availability

All data are available at https://github.com/tomlweeks1994/xxxx (full link to be added on acceptance)

## Code availability

The code to conduct analyses and replicate figures is available at https://github.com/tomlweeks1994/xxxx (full link to be added on acceptance)

## Extended Data Figures

**Extended Data Fig. 1.**
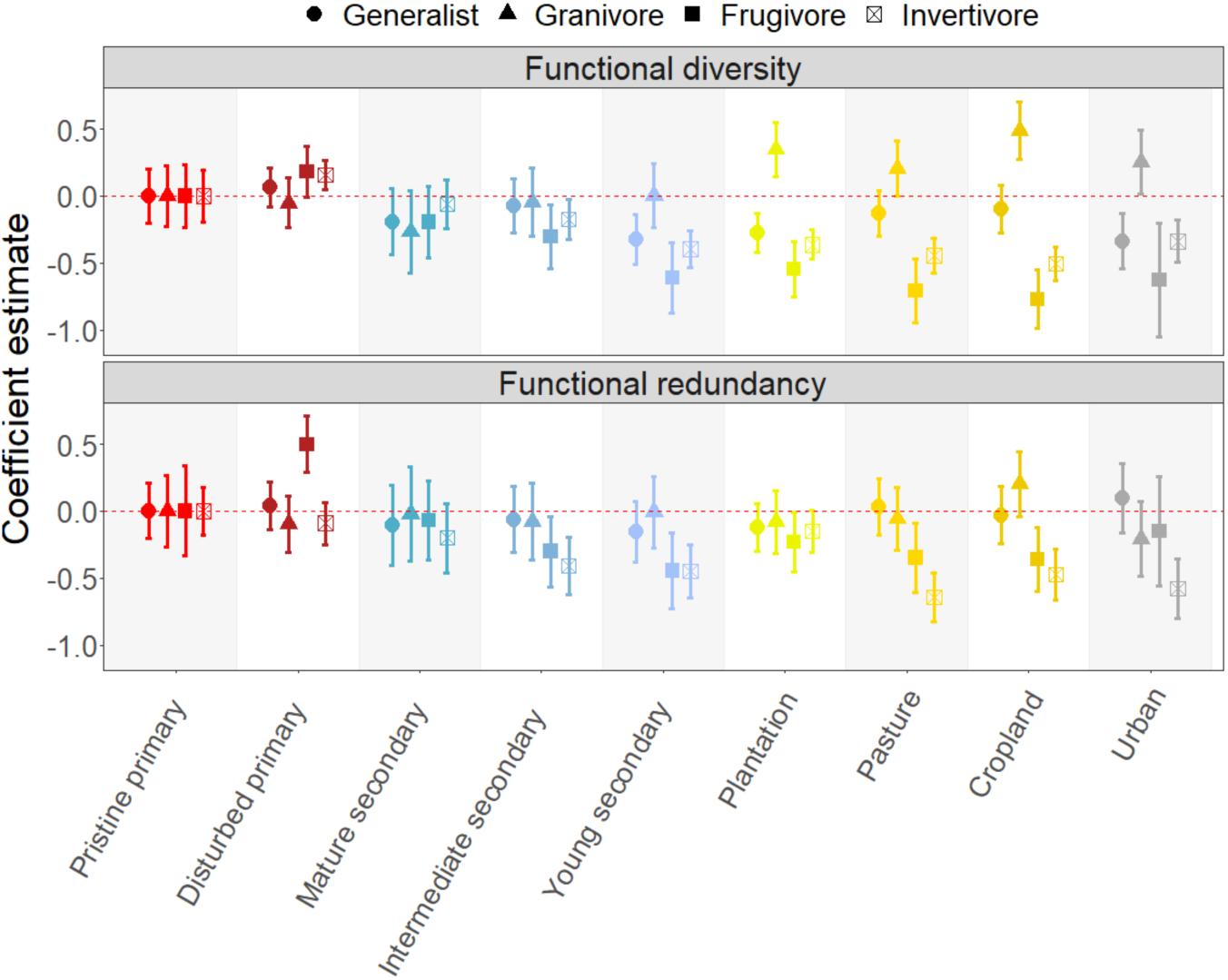
Land-use change alters the functionality and redundancy of avian assemblages. Results are outputs from univariate mixed effects models identifying how assemblage-level (**a**) Functional diversity (FD; measured as functional richness), and (**b**) Functional redundancy change in response to land-use. Metrics are calculated and compared across distinct functional guilds: generalists (*n* = 1256), granivores (*n* = 1051), frugivores (*n* = 617) & invertivores (*n* = 1264). Species are assigned to a functional guild if they consume greater than 60% of their diet from a single food source; all other species are defined as generalists. FD and functional redundancy values in pristine primary vegetation (including forests, grasslands and shrublands) are set to 0 for each dietary grouping. Functional metrics are then compared within group against this standardized pristine primary vegetation baseline value (dashed red line). Results shown are coefficient estimates and 95% confidence intervals. Both response variables were squareroot transformed prior to analysis and then scaled by their within-group standard deviation to aid comparison.

**Extended Data Fig. 2.**
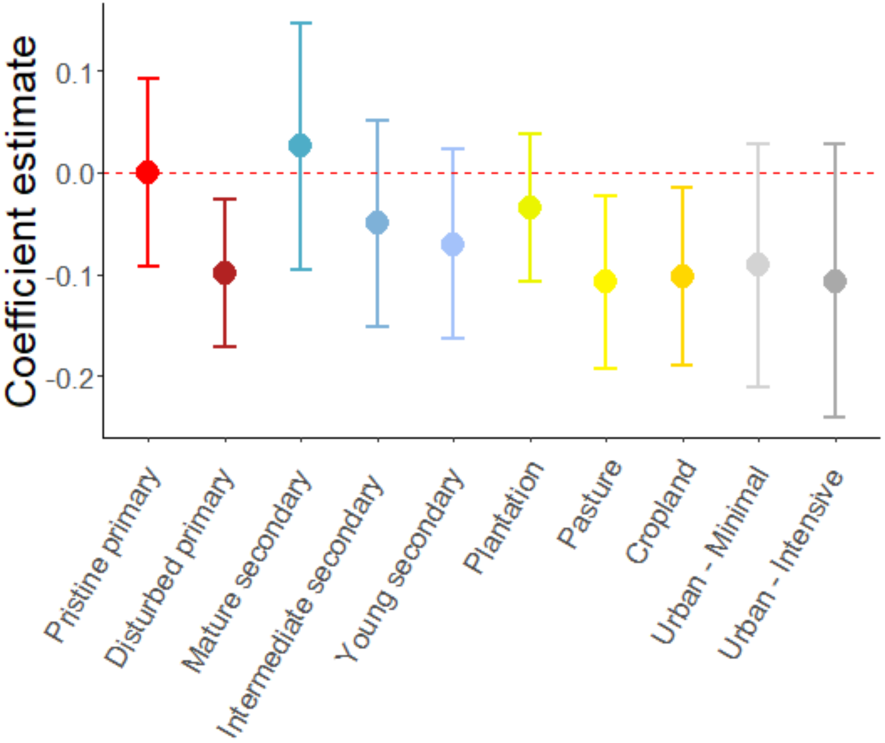
Effects of land-use change on rarity-based functional vulnerability. Results shown are the coefficient estimates and 95% confidence intervals derived from a univariate linear mixed-effects model assessing the relationship between land-use change and the rarity-based estimate of functional vulnerability (FV; *n* = 1278). For each species, rarity-based FV is calculated as the Pearson’s correlation coefficient between the species’ abundance and individual redundancy scores. Redundancy scores are calculated for each species as the change in assemblage-redundancy when the species is removed from the original assemblage. Decreasing coefficient estimates suggest that the most sensitive (rare) species are becoming relatively more redundant compared to pristine primary vegetation.

**Extended Data Fig. 3.**
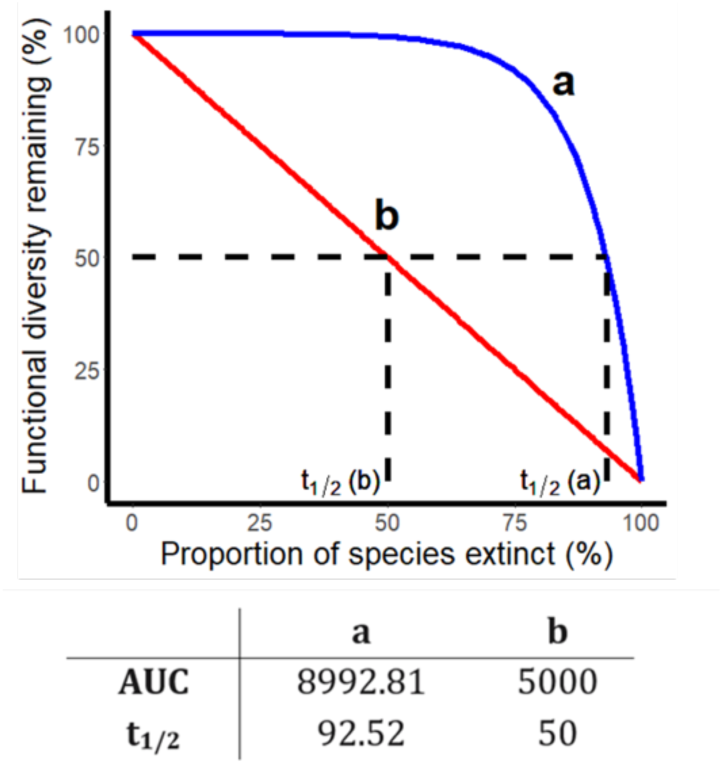
Theoretical representation of extinction curves. Coloured lines show two alternative types of extinction curves created by tracking the decline in functional diversity (FD; measured as functional richness) caused by the sequential removal of species from an assemblage. Blue extinction curve (**a**) depicts an assemblage with strong functional resistance (able to maintain relatively high FD despite increasing species loss). Red extinction curve (**b**) depicts an assemblage with weak functional resistance (FD declining linearly with species loss). We estimate functional resistance of the assemblage in two ways: the area under the curve (AUC), and the proportion of species remaining after the FD remaining is reduced to 50% (half-life; t_1/2_).

**Extended Data Fig. 4.**
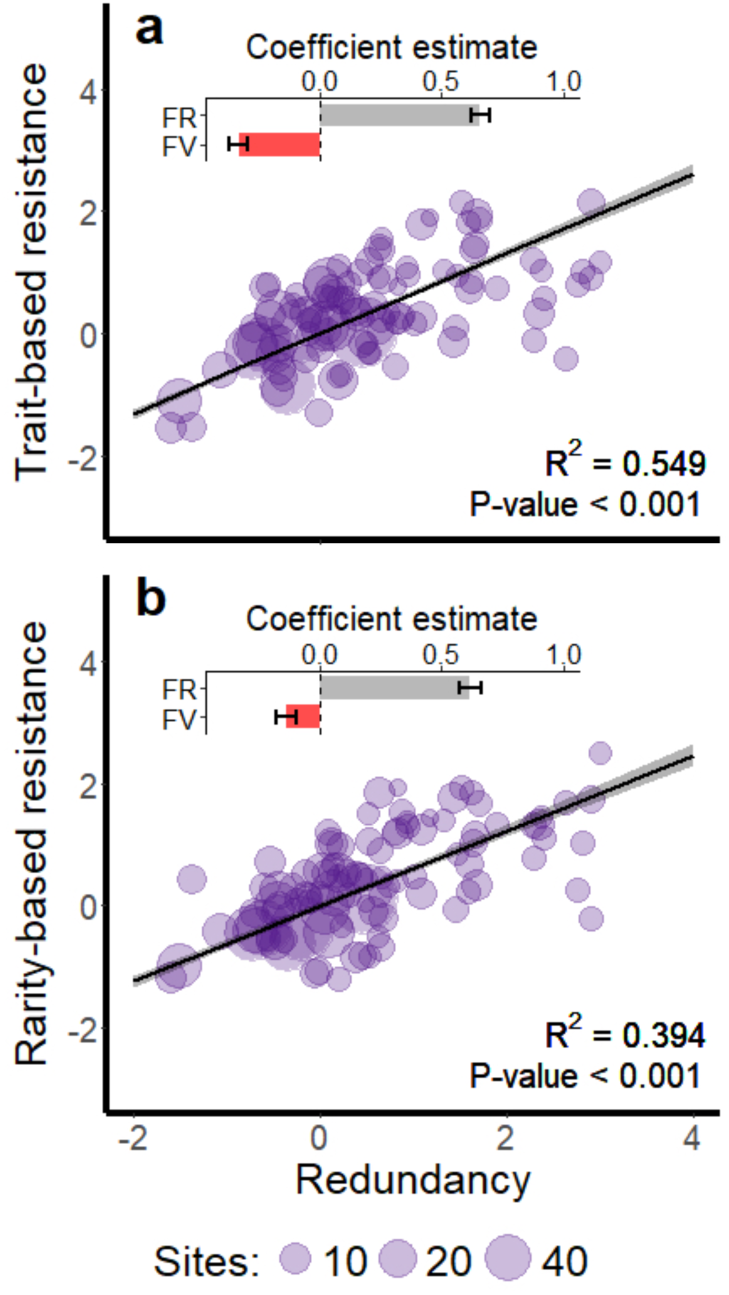
Proximal drivers of functional resistance to simulated species losses. Panels show the effect of functional redundancy (FR) and functional vulnerability (FV) on functional resistance. FR is the proportion of functional space shared by multiple species; FV is Pearson’s correlation coefficient between species-level redundancy and sensitivity scores; functional resistance values are calculated as the area under the extinction curve (AUC) for each assemblage (*n* = 1281) when species are removed under different species loss scenarios. Trait-based resistance (**a**) involved species removal in order of sensitivity to land-use change according to four response traits (geographical range size, body mass, dispersal ability and dietary specialism); rarity-based resistance (**b**) involved species removal in reverse order of abundance. Inset plots show results from a bivariate model analysing the effect of FR and FV on functional resistance across all assemblages (see Methods). To aid visualisation, points plotted are the average redundancy and resistance values for each study. The size of the points indicates the number of assemblages in each study. Solid black line is the slope of the coefficient estimate for redundancy; grey shaded area represents 95% confidence intervals.

**Extended Data Fig. 5.**
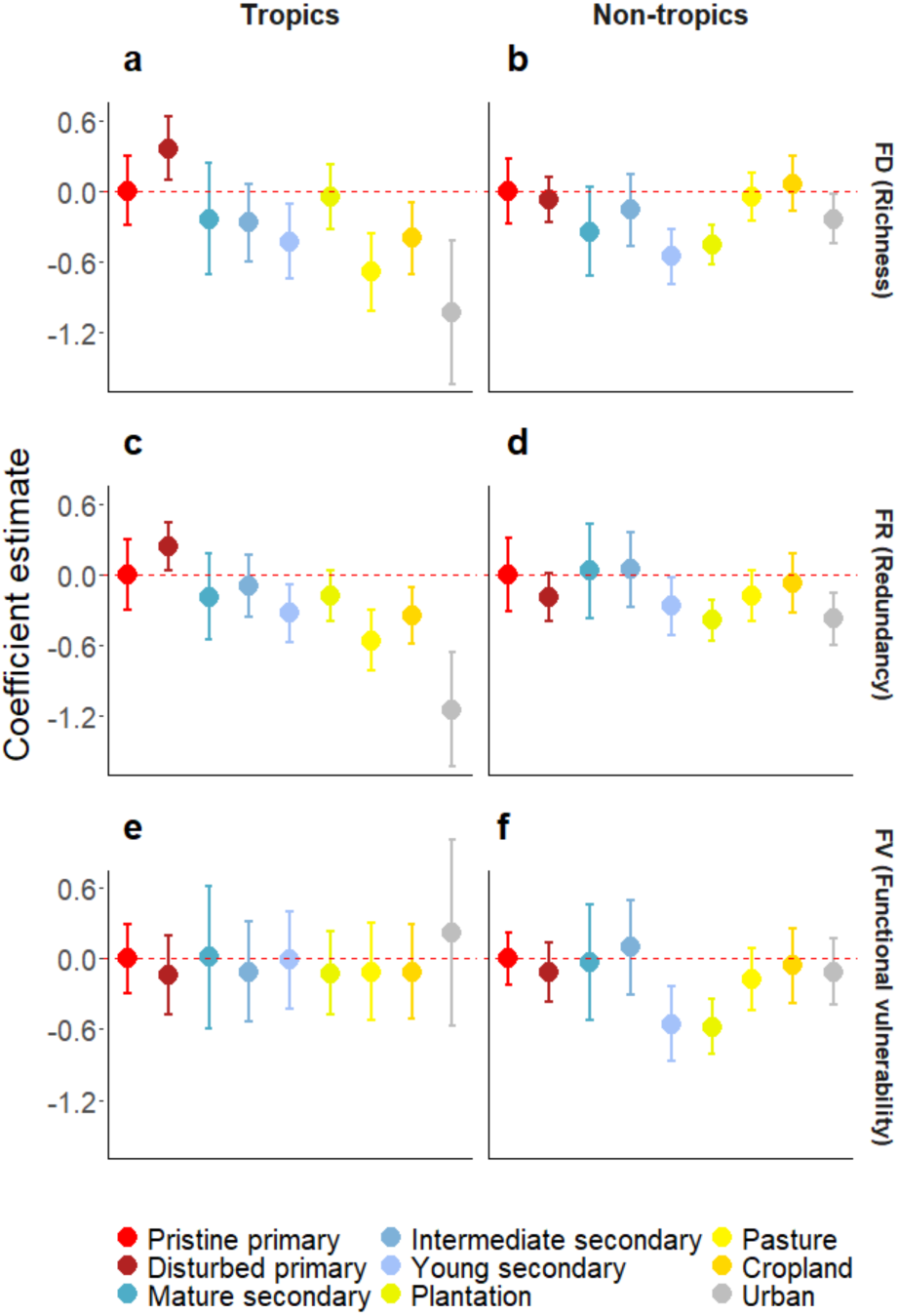
Impacts of land-use change on structure and function of avian assemblages vary by latitude. Results are outputs from univariate mixed effects models exploring effects of land-use change on (**a, b**) assemblage functional diversity (FD; measured as functional richness); (**c, d**) functional redundancy (FR), and (**e, f**) functional vulnerability (FV). Metrics are calculated and analyzed seperately for Tropical (*n* = 613), and Non-tropical (*n* = 668) sites. FV is calculated using Pearson’s correlation coefficient between species-level redundancy and trait-based sensitivity scores (see Methods). In all cases, metrics are compared to the baseline value for pristine primary vegation (forests, grasslands, shrublands and wetlands; dashed red line). Results shown are coefficient estimates and 95% confidence intervals. Response variables were squareroot transformed prior to analysis and then scaled by their standard deviation to aid comparison (the transformation step was not possible for FV).

**Extended Data Fig. 6.**
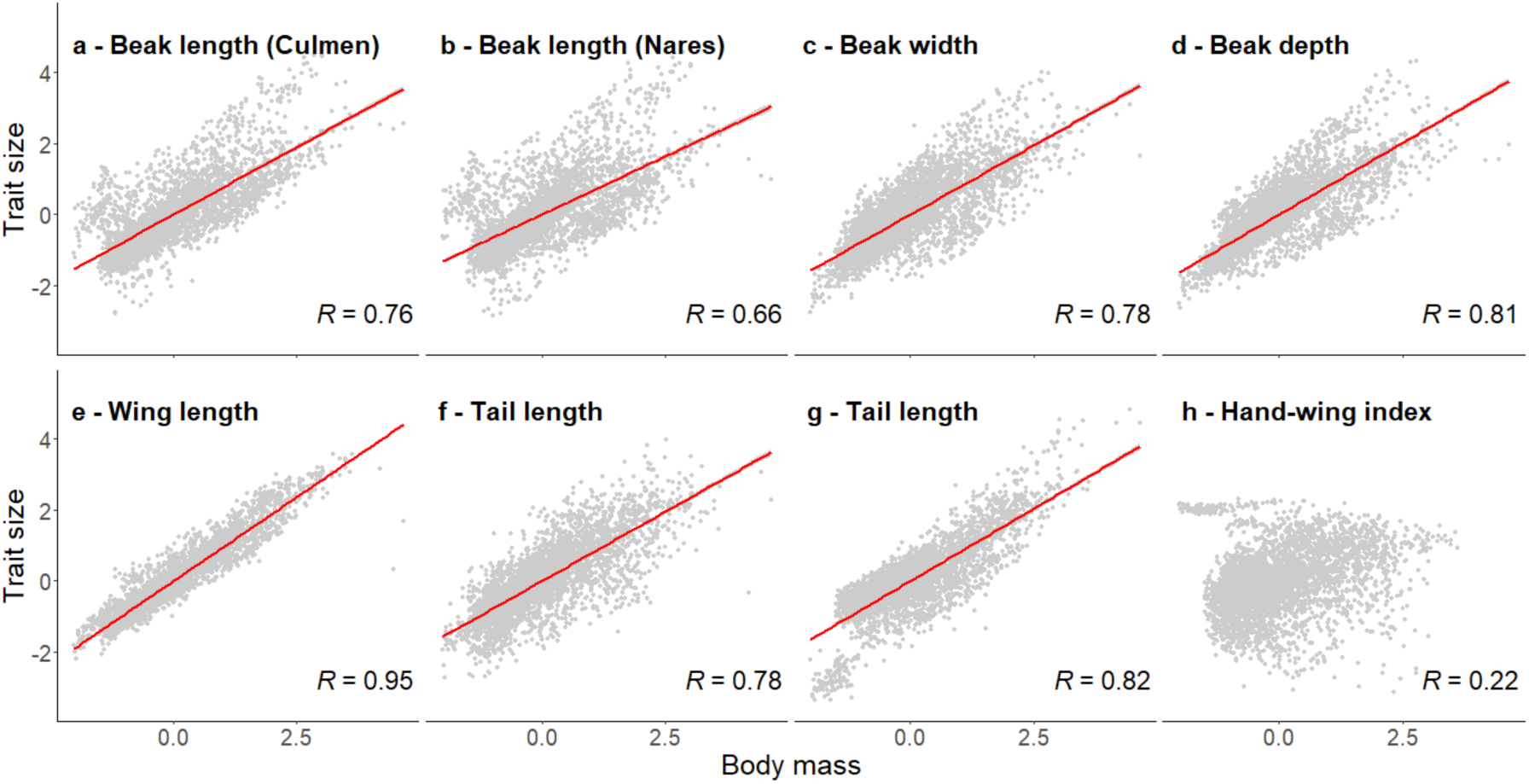
Correlations between morphological traits and body mass. Body mass has a strong positive correlation with most of our morphological trait values (**a-g**). Only hand wing index (**h**) is uncorrelated with body mass. Trait size and body mass values shown are logarithmically transformed and then scaled to their standard deviation. Red line indicates the slope of the correlation coefficient estimate. R-values are the Pearson’s correlation coefficients.

**Extended Data Fig. 7.**
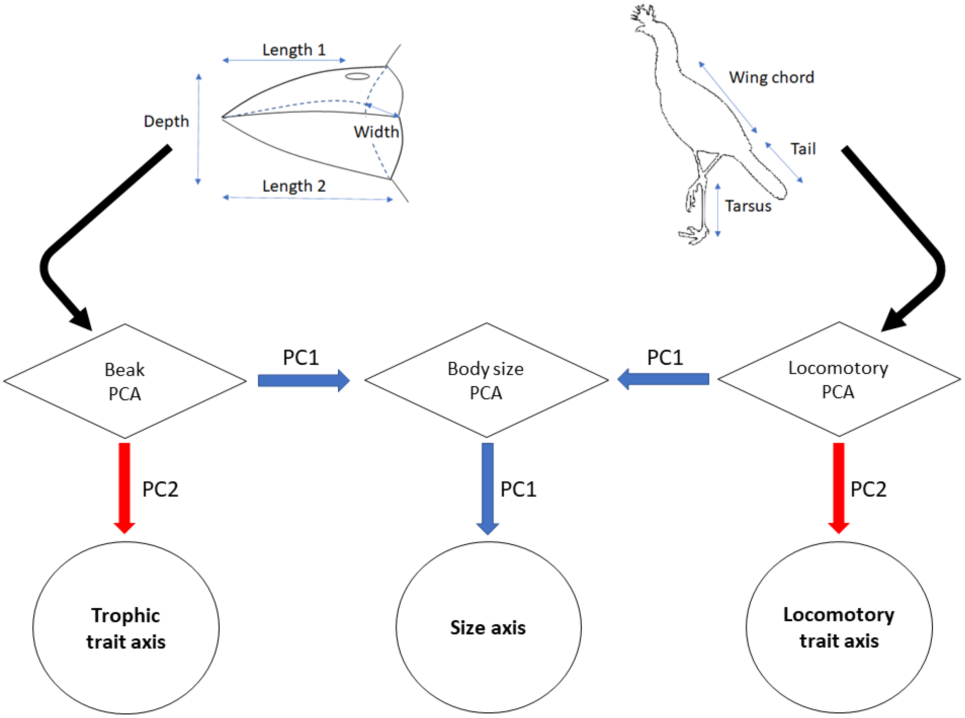
Using traits to quantify independent niche dimensions. Flow diagram illustrates the 2-step principal component analysis (PCA) used to generate three distinct trait axes. We separated seven morphological traits correlated with body mass into a trophic axis based on beak measurements (culmen length, tip-to-nares, depth, width) and a locomotory axis (wing length, tail length, tarsus length). We then performed two separate PCAs on trophic and locomotory traits using all species in the dataset. The first principal components (PCs) are correlated with overall size, so we used the second principal component to represent the dominant axis of variation, independent of body size, for beak shape and locomotion, respectively (Extended Data Fig. 7). After taking these second PCs as our trophic and locomotory trait axes, we performed a third PCA using both first principal components of the original PCAs and used the first principal component of this final PCA to reflect the size axis.

**Extended Data Fig. 8.**
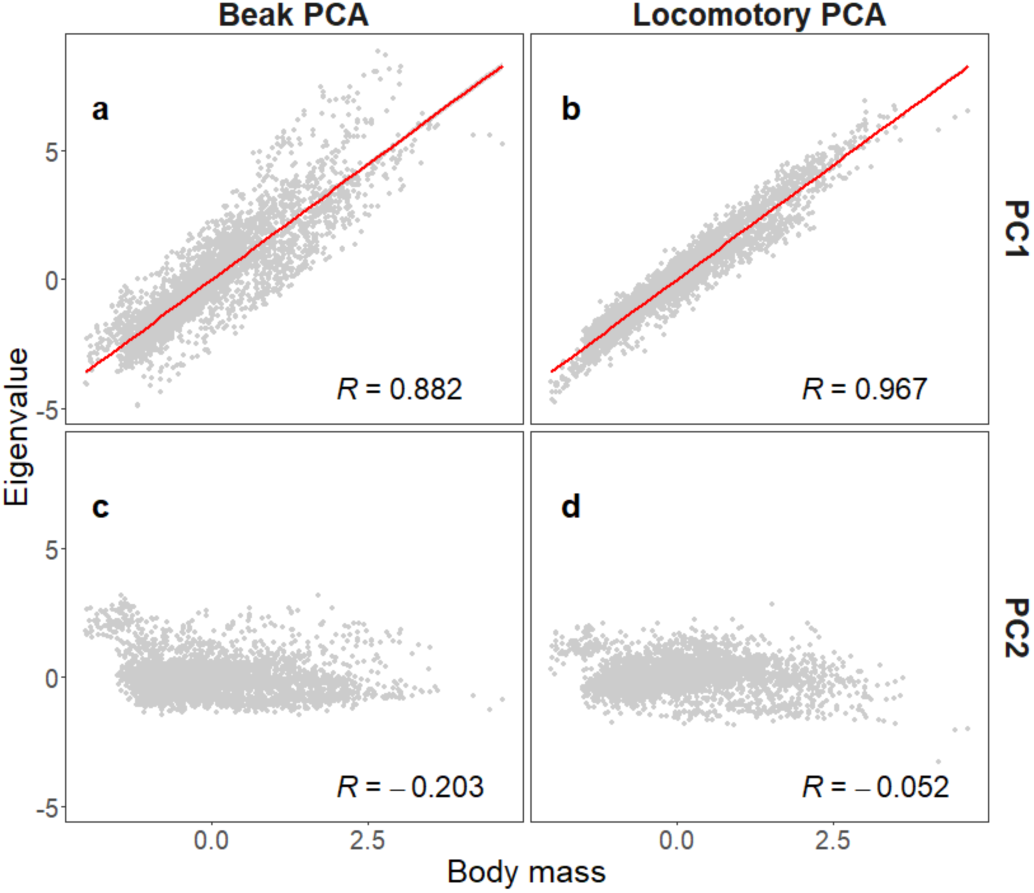
Correlations between principal component analysis outputs and body mass. To remove correlation between our trait values and overall species body size, we performed a 2-step principal component analysis (PCA) using two subsets of our trait-data. Beak-PCA was performed using four beak morphometrics: beak length (culmen), beak length (nares), beak width & beak depth. Locomotory-PCA was performed using three traits: wing length, tail length & tarsus length. The first principal components (**a, b**) of beak and locomotory trait PCAs are positively correlated with body mass. The second principal components (**c, d**) are uncorrelated with body mass. Body mass is logarithmically transformed and scaled to its standard deviation. Red line indicates the slope of the correlation coefficient estimate. R-values are Pearson’s correlation coefficients.

## References

Allan, E., Manning, P., Alt, F., Binkenstein, J., Blaser, S., Blüthgen, N., Böhm, S., Grassein, F., Hölzel, N., Klaus, V. H., Kleinebecker, T., Morris, E. K., Oelmann, Y., Prati, D., Renner, S. C., Rillig, M. C., Schaefer, M., Schloter, M., Schmitt, B., … Fischer, M. (2015). Land use intensification alters ecosystem multifunctionality via loss of biodiversity and changes to functional composition. Ecol. Lett. 18, 834–843.

Balmford, A. (1996). Extinction filters and current resilience: the significance of past selection pressures for conservation biology. Trends Ecol. Evol. 11, 193–196.

Barlow, J., Mestre, L. A. M., Gardner, T. A., & Peres, C. A. (2007). The value of primary, secondary and plantation forests for Amazonian birds. Biol. Conserv. 136, 212–231.

Betts, M. G., Wolf, C., Pfeifer, M., Banks-Leite, C., Arroyo-Rodríguez, V., Ribeiro, D. B., Barlow, J., Eigenbrod, F., Faria, D., Fletcher, R. J., Hadley, A. S., Hawes, J. E., Holt, R. D., Klingbeil, B., Kormann, U., Lens, L., Levi, T., Medina-Rangel, G. F., Melles, S. L., … Ewers, R. M. (2019). Extinction filters mediate the global effects of habitat fragmentation on animals. Science 366, 1236–1239.

Biggs, C. R., Yeager, L. A., Bolser, D. G., Bonsell, C., Dichiera, A. M., Hou, Z., Keyser, S. R., Khursigara, A. J., Lu, K., Muth, A. F., Negrete, B., & Erisman, B. E. (2020). Does functional redundancy affect ecological stability and resilience? A review and meta-analysis. Ecosphere 11, e03184.

BirdLife International. (2021). BirdLife Data Zone. Available online at: http://www.Birdlife.org/datazone [Accessed January 24, 2021].

Blonder, B., Morrow, C. B., Maitner, B., Harris, D. J., Lamanna, C., Violle, C., Enquist, B. J., & Kerkhoff, A. J. (2018). New approaches for delineating n-dimensional hypervolumes. *Meth*. Ecol. Evol. 9, 305–319.

Bregman, T. P., Lees, A. C., MacGregor, H. E. A., Darski, B., de Moura, N. G., Aleixo, A., Barlow, J., & Tobias, J. A. (2016). Using avian functional traits to assess the impact of land-cover change on ecosystem processes linked to resilience in tropical forests. Proc. R. Soc. B. 283, 20161289.

Bregman, T. P., Şekercioğlu, C. H., & Tobias, J. A. (2014). Global patterns and predictors of bird species responses to forest fragmentation: Implications for ecosystem function and conservation. Biol. Conserv. 169, 372–383.

Bright, J. A., Marugán-Lobón, J., Cobb, S. N., & Rayfield, E. J. (2016). The shapes of bird beaks are highly controlled by nondietary factors. Proc. Natl. Acad. Sci. USA 113, 5352–5357.

Burin, G., Kissling, W. D., Guimarães, P. R., Şekercioğlu, C. H., & Quental, T. B. (2016). Omnivory in birds is a macroevolutionary sink. Nat. Comm. 7, 11250.

Burkle, L. A., Marlin, J. C., & Knight, T. M. (2013). Plant-pollinator interactions over 120 years: loss of species, co-occurrence, and function. Science 339, 1611–1615.

Cadotte, M. W., Carscadden, K., & Mirotchnick, N. (2011). Beyond species: Functional diversity and the maintenance of ecological processes and services. J. Applied Ecol. 48, 1079–1087.

Carmona, C. P., de Bello, F., Mason, N. W. H., & Lepš, J. (2016). Traits without borders: integrating functional diversity across scales. Trends Ecol. Evol. 31, 382–394.

Carmona, C. P., de Bello, F., Mason, N. W. H., & Lepš, J. (2019). Trait probability density (TPD): measuring functional diversity across scales based on TPD with R. Ecology, 100 e02876.

Carmona, C. P., Tamme, R., Pärtel, M., De Bello, F., Brosse, S., Capdevila, P., González, R. M., González-Suárez, M., Salguero-Gómez, R., Vásquez-Valderrama, M., & Toussaint, A. (2021). Erosion of global functional diversity across the tree of life. Sci. Adv. 7, eabf2675.

Chapman, P. M., Tobias, J. A., Edwards, D. P., & Davies, R. G. (2018). Contrasting impacts of land-use change on phylogenetic and functional diversity of tropical forest birds. J. Applied Ecol. 55, 1604–1614.

Clapcott, J. E., Collier, K. J., Death, R. G., Goodwin, E. O., Harding, J. S., Kelly, D., … & Young, R. G. (2012). Quantifying relationships between land-use gradients and structural and functional indicators of stream ecological integrity. Freshw. Biol. 57, 74– 90.

Clavel, J., Julliard, R., & Devictor, V. (2010) Worldwide decline of specialist species: toward a global functional homogenization? Frontiers Ecol. Environ. 9, 222–228.

Croci, S., Butet, A., & Clergeau, P. (2008). Does urbanization filter birds on the basis of their biological traits. Condor 110, 223–240.

Cooke, R. S. C., Bates, A. E., & Eigenbrod, F. (2019). Global trade-offs of functional redundancy and functional dispersion for birds and mammals. Glob. Ecol. Biogeog. 28, 484–495.

Curtis, J. R., Robinson, W. D., Rompré, G., Moore, R. P., & McCune, B. (2021). Erosion of tropical bird diversity over a century is influenced by abundance, diet and subtle climatic tolerances. Sci. Rep. 11, 10045.

Crooks, K. R., & Soulé, M. E. (1999). Mesopredator release and avifaunal extinctions in a fragmented system. Nature 400, 563–566.

Davies, K. F., Margules, C. R., & Lawrence, J. F. (2004). A synergistic effect puts rare, specialized species at greater risk of extinction. Ecology 85, 265–271.

De Bello, F., Botta-Dukát, Z., Lepš, J., & Fibich, P. (2021). Towards a more balanced combination of multiple traits when computing functional differences between species. Meth. Ecol. Evol. 12, 443–448.

De Coster, G., Banks-Leite, C., & Metzger, J. P. (2015). Atlantic forest bird communities provide different but not fewer functions after habitat loss. Proc. R. Soc. B 282, 20142844.

De Vries, F. T., Liiri, M. E., Bjørnlund, L., Bowker, M. A., Christensen, S., Setälä, H. M., & Bardgett, R. D. (2012). Land use alters the resistance and resilience of soil food webs to drought. Nat. Clim. Chang. 2, 276–280.

Díaz, S., Purvis, A., Cornelissen, J. H. C., Mace, G. M., Donoghue, M. J., Ewers, R. M., Jordano, P., & Pearse, W. D. (2013). Functional traits, the phylogeny of function, and ecosystem service vulnerability. Ecol. Evol. 3, 2958–2975.

Drapeau, P., Leduc, A., Giroux, J. F., Savard, J. P. L., Bergeron, Y., & Vickery, W. L. (2000). Landscape-scale disturbances and changes in bird communities of boreal mixed-wood forests. Ecol. Monogr. 70, 423–444.

Duong, T. (2007). Ks: Kernel density estimation and kernel discriminant analysis for multivariate data in R. J. Stat. Softw. 21, 1–16.

Edwards, D. P., Tobias, J. A., Sheil, D., Meijaard, E., Laurance, W. F. (2014a). Maintaining ecosystem function and services in logged tropical forests. Trends Ecol. Evol. 29, 511– 520.

Edwards, D. P., Magrach, A., Woodcock, P., Ji, Y., Lim, N. T. L., Edwards, F. A., Larsen, T. H., Hsu, W. W., Benedick, S., Khen, C. V., Chung, A. Y. C., Reynolds, G., Fisher, B., Laurance, W. F., Wilcove, D. S., Hamer, K. C., & Yu, D. W. (2014b). Selective-logging and oil palm: Multitaxon impacts, biodiversity indicators, and trade-offs for conservation planning. Ecol. Appl. 24, 2029–2049.

Edwards, F. A., Edwards, D. P., Larsen, T. H., Hsu, W. W., Benedick, S., Chung, A., … & Hamer, K. C. (2013). Does logging and forest conversion to oil palm agriculture alter functional diversity in a biodiversity hotspot? Anim. Conserv. 17, 163–173.

Etard, A., Pigot, A. L., & Newbold, T. (2022). Intensive human land uses negatively affect vertebrate functional diversity. Ecol. Lett. 25, 330–343.

Felice, R. N., Tobias, J. A., Pigot, A. L., & Goswami, A. (2019). Dietary niche and the evolution of cranial morphology in birds. Proc. R. Soc. B 286, 20182677.

Fischer, J., Lindenmayer, D. B., Blomberg, S. P., Montague-Drake, R., Felton, A., & Stein, J. A. (2007). Functional richness and relative resilience of bird communities in regions with different land use intensities. Ecosystems 10, 964–974.

Flynn, D. F. B., Gogol-Prokurat, M., Nogeire, T., Molinari, N., Richers, B. T., Lin, B. B., Simpson, N., Mayfield, M. M., & DeClerck, F. (2009). Loss of functional diversity under land use intensification across multiple taxa. Ecol. Lett. 12, 22–33.

Foley, J. A., DeFries, R., Asner, G. P., Barford, C., Bonan, G., Carpenter, S. R., Chapin, F. S., Coe, M. T., Daily, G. C., Gibbs, H. K., Helkowski, J. H., Holloway, T., Howard, E. A., Kucharik, C. J., Monfreda, C., Patz, J. A., Prentice, I. C., Ramankutty, N., & Snyder, P. K. (2005). Global consequences of land use. Science 309, 570–574.

Fonseca, C. R., & Ganade, G. (2001). Species functional redundancy, random extinctions and the stability of ecosystems. J. Ecol. 89, 118–125.

Fraixedas, S., Lindén, A., Piha, M., Cabeza, M., Gregory, R., & Lehikoinen, A. (2020). A state-of-the-art review on birds as indicators of biodiversity: Advances, challenges, and future directions. Ecol. Indic. 118, 106728.

Galland, T., Pérez Carmona, C., Götzenberger, L., Valencia, E., & de Bello, F. (2020). Are redundancy indices redundant? An evaluation based on parameterized simulations. Ecol. Indic. 116, 106488.

Gaston, K. J., & Blackburn, T. M. (1995). Birds, body size and the threat of extinction. Phil. Trans. R. Soc. B 347, 205–212.

Gorczynski, D., & Beaudrot, L. (2021). Functional diversity and redundancy of tropical forest mammals over time. Biotropica 53, 51–62.

Grimm, N. B., Faeth, S. H., Golubiewski, N. E., Redman, C. L., Wu, J., Bai, X., & Briggs, J. M. (2008). Global change and the ecology of cities. Science 319, 756–760.

Hillebrand, H., Langenheder, S., Lebret, K., Lindström, E., Östman, Ö., & Striebel, M. (2018). Decomposing multiple dimensions of stability in global change experiments. Ecol. Lett. 21, 21–30.

Hudson, L. N., Newbold, T., Contu, S., Hill, S. L. L., Lysenko, I., De Palma, A., Phillips, H. R. P., Senior, R. A., Bennett, D. J., Booth, H., Choimes, A., Correia, D. L. P., Day, J., Echeverría-Londoño, S., Garon, M., Harrison, M. L. K., Ingram, D. J., Jung, M., Kemp, V., … Purvis, A. (2014). The PREDICTS database: A global database of how local terrestrial biodiversity responds to human impacts. Ecol. Evol. 4, 4701–4735.

Hughes, A. C., Orr, M. C., Ma, K., Costello, M. J., Waller, J., Provoost, P., … & Qiao, H. (2021). Sampling biases shape our view of the natural world. Ecography 44, 1259–1269.

Jaureguiberry, P. et al. (2022) The direct drivers of recent global anthropogenic biodiversity loss. Sci. Adv. 8, eabm9982.

La Sorte, F. A., Lepczyk, C. A., Aronson, M. F. J., Goddard, M. A., Hedblom, M., Katti, M., MacGregor-Fors, I., Mörtberg, U., Nilon, C. H., Warren, P. S., Williams, N. S. G. & Yang, J. (2018). The phylogenetic and functional diversity of regional breeding bird assemblages is reduced and constricted through urbanization. Divers. Distrib. 24, 928– 938.

Laliberté, E., Wells, J. A., DeClerck, F., Metcalfe, D. J., Catterall, C. P., Queiroz, C., Aubin, I., Bonser, S. P., Ding, Y., Fraterrigo, J. M., McNamara, S., Morgan, J. W., Merlos, D. S., Vesk, P. A. & Mayfield, M. M. (2010). Land-use intensification reduces functional redundancy and response diversity in plant communities. Ecol. Lett. 13, 76–86.

Lande, R. (1993). Risks of population extinction from demographic and environmental stochasticity and random catastrophes. Amer. Nat. 142, 911–927.

Lawton, J. H., & Brown, V. K. (1994). *Redundancy in ecosystems*. In Biodiversity and ecosystem function (pp. 255–270). Springer, Berlin, Heidelberg.

Leclère, D., Obersteiner, M., Barrett, M., Butchart, S. H., Chaudhary, A., De Palma, A., … & Young, L. (2020). Bending the curve of terrestrial biodiversity needs an integrated strategy. Nature 585, 551–556.

Legendre, P., & Legendre, L. (1998). Numerical Ecology. Elsevier, Amsterdam.

Leitão, R. P., Zuanon, J., Villéger, S., Williams, S. E., Baraloto, C., Fortune, C., Mendonça, F. P., & Mouillot, D. (2016). Rare species contribute disproportionately to the functional structure of species assemblages. Proc. R. Soc. B 283, 20160084.

Loreau, M, Naeem, S., Inchausti, P., Bengtsson, J., Grime, J. P., Hector, A., Hooper, D. U., Huston, M. A., Raffaelli, D., Schmid, B., Tilman, D., & Wardle, D. A. (2001). Biodiversity and ecosystem functioning: Current knowledge and future challenges. Science 294, 804–808.

Luck, G. W., Carter, A., & Smallbone, L. (2013). Changes in bird functional diversity across multiple land uses: Interpretations of functional redundancy depend on functional group identity. PloS ONE 8, e63671.

Magnago, L.F.S., Edwards, D.P., Edwards, F.A., Magrach, A., Martins, S.V., & Laurance, W.F. (2014) Functional attributes change but functional richness is unchanged after fragmentation of Brazilian Atlantic forests. J. Ecol. 102, 475–485.

McCann, K. S. (2000). The diversity–stability debate. Nature 405, 228–233.

McLean, M., Auber, A., Graham, N. A., Houk, P., Villéger, S., Violle, C., … & Mouillot, D. (2019). Trait structure and redundancy determine sensitivity to disturbance in marine fish communities. Glob. Change Biol. 25, 3424–3437.

McKinney, M. L., & Lockwood, J. L. (1999). Biotic homogenization: A few winners replacing many losers in the next mass extinction. Trends Ecol. Evol. 14, 450–453.

Mouillot, D., Bellwood, D. R., Baraloto, C., Chave, J., Galzin, R., Harmelin-Vivien, M., … & Thuiller, W. (2013). Rare species support vulnerable functions in high-diversity ecosystems. PloS Biol. 11, e1001569.

Mouillot, D., Villéger, S., Parravicini, V., Kulbicki, M., Arias-González, J. E., Bender, M., … & Bellwood, D. R. (2014). Functional over-redundancy and high functional vulnerability in global fish faunas on tropical reefs. Proc. Natl. Acad. Sci. USA 111, 13757–13762.

Naeem, S. (1998). Species redundancy and ecosystem reliability. Conserv. Biol., 12, 39–45.

Naeem, S., & Li, S. (1997). Biodiversity enhances ecosystem reliability. Nature 390, 507–509.

Newbold, T., Scharlemann, J. P. W., Butchart, S. H. M., Şekercioǧlu, Ç. H., Alkemade, R., Booth, H., & Purves, D. W. (2013). Ecological traits affect the response of tropical forest bird species to land-use intensity. Proc. R. Soc. B 280, 20122131.

Newbold, T., Scharlemann, J. P. W., Butchart, S. H. M., Şekercioğlu, Ç. H., Joppa, L., Alkemade, R., & Purves, D. W. (2014). Functional traits, land-use change and the structure of present and future bird communities in tropical forests. Glob. Ecol. Biogeog. 23, 1073–1084.

Newbold, T., Hudson, L. N., Hill, S. L., Contu, S., Lysenko, I., Senior, R. A., … & Purvis, A. (2015). Global effects of land use on local terrestrial biodiversity. Nature 520, 45–50.

Newbold, T., Bentley, L. F., Hill, S. L. L., Edgar, M. J., Horton, M., Su, G., Şekercioğlu, Ç. H., Collen, B., & Purvis, A. (2020). Global effects of land use on biodiversity differ among functional groups. Funct. Ecol. 34, 684–693.

Nunes, C. A., Barlow, J., França, F., Berenguer, E., Solar, R. R. C., Louzada, J., Leitão, R., Maia, L., Oliveira, V. H. F., Braga, R. F., Vaz-de-Mello, F. Z., & Sayer, E. J. (2021). Functional redundancy of Amazonian dung beetles confers community-level resistance to primary forest disturbance. Biotropica 53, 1510–1521.

Nunes, C. A., Berenguer, E., França, F., Ferreira, J., Lees, A. C., Louzada, J., … & Barlow, J. (2022). Linking land-use and land-cover transitions to their ecological impact in the Amazon. Proc. Natl. Acad. Sci. USA 119, e2202310119.

Oliveira Hagen, E., Hagen, O., Ibáñez-Álamo, J. D., Petchey, O. L., & Evans, K. L. (2017). Impacts of urban areas and their characteristics on avian functional diversity. Front. Ecol. Evol. 5, 84.

Oliver, T. H., Heard, M. S., Isaac, N. J. B., Roy, D. B., Procter, D., Eigenbrod, F., Freckleton, R., Hector, A., Orme, C. D. L., Petchey, O. L., Proença, V., Raffaelli, D., Suttle, K. B., Mace, G. M., Martín-López, B., Woodcock, B. A., & Bullock, J. M. (2015). Biodiversity and resilience of ecosystem functions. Trends Ecol. Evol. 30, 673–684.

Olson, D. M., Dinerstein, E., Wikramanayake, E. D., Burgess, N. D., Powell, G. V. N., Underwood, E. C., D’Amico, J. A., Itoua, I., Strand, H. E., Morrison, J. C., Loucks, C. J., Allnutt, T. F., Ricketts, T. H., Kura, Y., Lamoreux, J. F., Wettengel, W. W., Hedao, P., Kassem, K. R. 2001. Terrestrial ecoregions of the world: a new map of life on Earth. Bioscience 51, 933–938.

Owens, I. P. F., & Bennett, P. M. (2000). Ecological basis of extinction risk in birds: Habitat loss versus human persecution and introduced predators. Proc. Natl. Acad. Sci. USA 97, 12144–12148.

Pakeman, R. J. (2014). Functional trait metrics are sensitive to the completeness of the species’ trait data? *Meth*. Ecol. Evol. 5, 9–15.

Petchey, O. L., Hector, A., & Gaston, K. J. (2004). How do different measures of functional diversity perform. Ecology 85, 847–857.

Pigot, A. L., Sheard, C., Miller, E. T., Bregman, T. P., Freeman, B. G., Roll, U., Seddon, N., Trisos, C. H., Weeks, B. C., & Tobias, J. A. (2020). Macroevolutionary convergence connects morphological form to ecological function in birds. *Nat*. Ecol. Evol. 4, 230– 239.

Sasaki, T., Yoshihara, Y., Takahashi, M., Byambatsetseg, L., Futahashi, R., Nyambayar, D., & Suyama, Y. (2017). Differential responses and mechanisms of productivity following experimental species loss scenarios. Oecologia 183, 785–795.

Şekercioğlu, Ç. H., Ehrlich, P. R., Daily, G. C., Aygen, D., Goehring, D., & Sandí, R. F. (2002). Disappearance of insectivorous birds from tropical forest fragments. Proc. Natl. Acad. Sci. USA 99, 263–267.

Şekercioğlu, Ç. H., Daily, G. C., & Ehrlich, P. R. (2004). Ecosystem consequences of bird declines. Proc. Natl. Acad. Sci. USA 101, 18042–18047.

Şekercioğlu, C. H. (2006). Increasing awareness of avian ecological function. Trends Ecol. Evol. 21, 464–471.

Sheard, C., Neate-Clegg, M. H. C., Alioravainen, N., Jones, S. E. I., Vincent, C., MacGregor, H. E. A., Bregman, T. P., Claramunt, S., & Tobias, J. A. (2020). Ecological drivers of global gradients in avian dispersal inferred from wing morphology. Nat. Comm. 11, 2463.

Sol, D., Trisos, C., Múrria, C., Jeliazkov, A., González-Lagos, C., Pigot, A. L., Ricotta, C., Swan, C. M., Tobias, J. A., & Pavoine, S. (2020). The worldwide impact of urbanisation on avian functional diversity. Ecol. Lett. 23, 962–972.

Suding, K. N., Lavorel, S., Chapin, F. S., Cornelissen, J. H. C., Diaz, S., Garnier, E., Goldberg, D., Hooper, D. U., Jackson, S. T. and Navas, M.-L. (2008) Scaling environmental change through the community-level: a trait-based response-and-effect framework for plants. Glob. Change Biol. 14, 1125–1140.

Tilman, D., Knops, J., Wedin, D., Reich, P., Ritchie, M., & Siemann, E. (1997). The influence of functional diversity and composition on ecosystem processes. Science 277, 1300–1302.

Tilman, D., Reich, P. B., & Knops, J. M. (2006). Biodiversity and ecosystem stability in a decade-long grassland experiment. Nature 441, 629–632.

Tobias, J. A., Sheard, C., Pigot, A. L., Devenish, A. J. M., Yang, J., Sayol, F., Neate-Clegg, M. H. C., Alioravainen, N., Weeks, T. L., Barber, R. A., Walkden, P. A., MacGregor, H. E. A., Jones, S. E. I., Vincent, C., Phillips, A. G., Marples, N. M., Montaño-Centellas, F. A., Leandro-Silva, V., Claramunt, S., … Schleuning, M. (2022). AVONET: morphological, ecological and geographical data for all birds. Ecol. Lett. 25, 581–597.

Trisos, C. H., Petchey, O. L., & Tobias, J. A. (2014). Unraveling the interplay of community assembly processes acting on multiple niche axes across spatial scales. Am. Nat. 184, 593–608.

Vandewalle, M., de Bello, F., Berg, M. P., Bolger, T., Dolédec, S., Dubs, F., Feld, C. K., Harrington, R., Harrison, P. A., Lavorel, S., da Silva, P. M., Moretti, M., Niemelä, J., Santos, P., Sattler, T., Sousa, J. P., Sykes, M. T., Vanbergen, A. J., & Woodcock, B. A. (2010). Functional traits as indicators of biodiversity response to land use changes across ecosystems and organisms. Biodivers. Conserv. 19, 2921–2947.

Villéger, S., Mason, N. W. H., & Mouillot, D. (2008). New multidimensional functional diversity indices for a multifaceted framework in functional ecology. Ecology 89, 2290– 2301.

Weeks, B. C., Naeem, S., Lasky, J. R. & Tobias, J. A. (2022a). Diversity and extinction risk are inversely related at a global scale. Ecol. Lett. 25, 697–707.

Weeks, B. C., O’Brien, B. K., Chu, J. J., Claramunt, S., Sheard, C., & Tobias, J. A. (2022b). Morphological adaptations linked to flight efficiency and aerial lifestyle determine natal dispersal distance in birds. Funct. Ecol. 36, 1681–1689.

Weeks, T.L., Betts, M.G., Pfeifer, M., Wolf, C., Banks-Leite, C., Barbaro, L., Barlow, J., Cerezo, A., Kennedy, C.M., Kormann, U.G., Marsh, C.J., Olivier, P.I., Phalan, B.T., Possingham, H.P., Wood, E.M., & Tobias, J.A. (2023) Climate-driven variation in dispersal ability predicts responses to forest fragmentation in birds. *Nat*. Ecol. Evol. 7, 1079–1091.

Whelan, C. J., Şekercioğlu, Ç. H., & Wenny, D. G. (2015). Why birds matter: from economic ornithology to ecosystem services. J. Ornithol. 156, 227–238.

White, A. E. (2016). Geographical barriers and dispersal propensity interact to limit range expansions of himalayan birds. Am. Nat. 188, 99–112.

Wilman, H., Belmaker, J., Simpson, J., de la Rosa, C., Rivadeneira, M. M., & Jetz, W. (2014). EltonTraits 1.0: Species-level foraging attributes of the world’s birds and mammals. Ecology 95, 2027.

Wunderle, J. M., Willig, M. R., & Henriques, L. M. P. (2005). Avian distribution in treefall gaps and understorey of terra firme forest in the lowland Amazon. Ibis 147, 109–129.

Zhang, S., & Zang, R. (2021). Tropical forests are vulnerable in terms of functional redundancy. Biol. Conserv. 262, 109326.

